# Vertical stratification of environmental DNA in the open ocean captures ecological patterns and behavior of deep-sea fishes

**DOI:** 10.1101/2021.02.10.430594

**Authors:** Oriol Canals, Iñaki Mendibil, María Santos, Xabier Irigoien, Naiara Rodríguez-Ezpeleta

## Abstract

The deep-sea remains among the most unknown ecosystems on Earth despite its relevant role in carbon sequestration and increasing threat due to interest by fishing and mining industries. This, together with the recent discovery that the upper layer of this ecosystem (mesopelagic zone) harbors about 90% of the fish biomass on Earth, claims for a deeper understanding of the deep-sea so that the foundations for a sustainable use of its resources can be established. The analysis of environmental DNA (eDNA) collected from the water column emerges as an alternative to traditional methods to acquire this elusive information, but its application to the deep ocean is still incipient. Here, we have amplified and sequenced the fish eDNA contained in vertical profile samples (from surface to 2000 m depth) collected during day and night-time throughout the Bay of Biscay. We found that eDNA-derived deep-sea fish richness and abundance follow a day-night pattern that is consistent with the diel migratory behavior of many mesopelagic species, and that eDNA can reveal species-specific distribution and movement through the water column. These results highlight the potential of eDNA-based studies to improve our knowledge on the species inhabiting the dark ocean before this still pristine ecosystem is exploited.

## Introduction

The deep-sea, i.e., the dark oceanic zone below 200 m depth, is the largest, least explored, and most pristine ecosystem on Earth (Sutton 2013; Duarte et al. 2020; Martin et al. 2020). Recent estimates suggest that the upper deep-sea layer (from 200 to 1000 m depth), referred to as the mesopelagic zone, may contain 90% of the total fish biomass (Irigoien et al. 2014), which, in the current context of increasing global human population and need to end overfishing, uncovers an alternative source of food for direct human consumption or as feed supply for aquaculture (FAO 2018). Yet, uncontrolled exploitation of this resource could result in irremediable damage (Mengerink et al. 2014; Martin et al. 2020) since mesopelagic fish play an important role in carbon sequestration (Davison et al. 2013; Thurber et al. 2014; Klevjer et al. 2016), and in trophic connectivity, by linking upper and lower ocean layers (St. John et al. 2016). This is due to the diel vertical migration (DVM) of many mesopelagic fishes moving into upper layers for feeding at nighttime and into deeper layers to avoid predation at daytime (Sutton 2013), which has been referred to as the “largest daily animal migration on Earth” (St. John et al. 2016).

Understanding the role of mesopelagic organisms in global biogeochemical cycles and evaluating the ecosystem services they provide is urgent before uninformed extraction of these resources starts (Martin et al. 2020). Fundamental knowledge gaps need to be filled for this purpose, such as which animals inhabit the mesopelagic zone, how are they distributed in space and time and which factors control their distribution. This lack of information is mostly due to the inaccessibility of deep-sea ecosystems, challenging traditional sampling methods (Martin et al. 2020). Although net tows, video plankton recorders, bioacoustics, and autonomous or remotely operated underwater vehicles are being applied, they are either extremely time-consuming, expensive, selective, highly specialized, hardly available for most research groups, and/or reliant on often scarce taxonomic expertise (Hansen et al. 2018). Thus, new technologies are required to acquire the needed insights on deep-sea ecosystems at a global scale to set the basics for studies aiming at evaluating the role of mesopelagic organisms as diet of commercially important and emblematic species, at understanding their role in carbon sequestration, and at exploring their potential as food and bioactive compound sources.

In this scenario, environmental DNA (eDNA) based approaches arise as cost-effective alternatives to explore the deep-sea. Water column environmental DNA includes DNA traces released by macro-organisms (including fish) into the environment in form of tissue, cells, scales, mucus, or feces. This material can be analyzed through metabarcoding, which allows community characterization by simultaneously sequencing a short region of the genome of multiple species, which is then compared to a reference database. eDNA collected from water samples can provide biodiversity information of virtually any marine habitat (Thomsen et al. 2012) including deep-sea environments (Thomsen et al. 2016; Laroche et al. 2020; McClenaghan et al. 2020). However, inferring the macro-organisms present in marine ecosystems by analyzing their DNA released into the environment requires an understanding of how DNA traces behave in the water column, and many factors related to extra-organismal DNA production rate, degradation and transport have been reported to influence its detection and concentration in the ocean (Hansen et al. 2018).

To date and to the best of our knowledge, no study has assessed whether eDNA is vertically structured or mixed up along the water column in the open ocean, which is crucial to evaluate the potential of eDNA to assess diversity and behavior of deep-sea fish communities. Here, we have applied eDNA metabarcoding to eight vertical profiles (up to 2000m depth) collected during day and night-time along the continental slope of the Bay of Biscay. We hypothesized that fish eDNA is vertically stratified along the oceanic water column, and can therefore provide information on fish vertical structuring. Evidence of eDNA-based approaches being capable of inferring deep-sea fish ecology, including diversity and vertical distribution, will greatly broaden the horizon of exploring new approaches for further understanding the remote deep-sea, which is key to ensure a sustainable exploitation of its resources.

## Material and Methods

### Sampling

Vertical profile samples from eight stations along the continental slope of the Bay of Biscay (Figure 1; Supplementary Table S1) were collected in Spring 2018 on board the Ramón Margalef research vessel at 5, 50, 200, 500 and 1,000m depth and at 50 m above the seafloor (thereafter > 1,000 m) using a rosette sampler and, from five of the stations, also at 4.4 m depth using the continuous clean circuit intake of the ship. On board, 2-5 liters of 50 μm pore-sized mesh prefiltered water samples were immediately filtered through Sterivex 0.45 μm pore size enclosed filters (Millipore) using a peristaltic pump and kept at −20 °C until further processing. In total, 52 samples from 8 vertical profiles (13 from the surface and 8 from each 50, 200, 500, 1,000 and >1,000 m depth) were collected.

**Figure 1.**
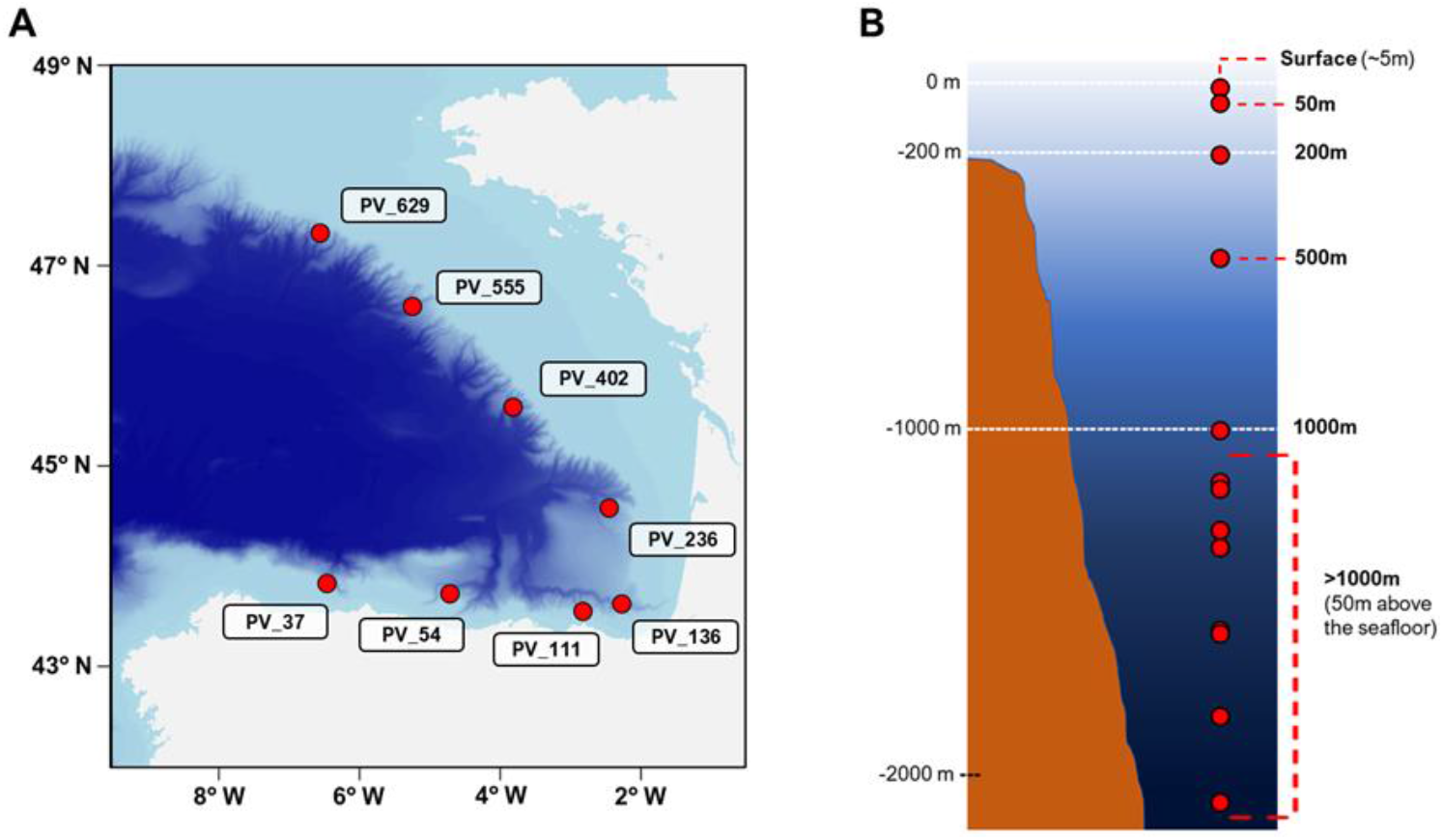
**A:** Map of Bay of Biscay displaying the eight stations from where samples used in this study were collected. **B:** Depths at which samples were collected in each of the eight vertical profiles.

### DNA extraction and metabarcoding

Laboratory and bioinformatic analysis procedures for eDNA metabarcoding were performed as described in Fraija-Fernández et al. (2020) and are detailed in Supplementary Information. Briefly, a short region of the 12S ribosomal RNA gene (Valentini et al. 2016) was amplified from the extracted eDNA and sequenced. Resulting sequences were quality filtered and taxonomically assigned using an in-house curated database restricted to the fish species (Myxini, Petromyzonti, Holocephali, Elasmobranchii, Sarcopterygii and Actinopterygii) expected in the Northeast Atlantic and Mediterranean areas. Raw reads from duplicate surface samples (those collected by the rosette and the intake circuit) were pooled into one sample. Only reads assigned to the species and genus levels were considered for subsequent analyses, and those reads assigned to species of the same genus were grouped. Deep-sea fish were classified as such for spending most of their lifetime below 200 m depth based on FishBase (http://www.fishbase.org/).

## Results

### Fish diversity in the continental slope of the Bay of Biscay

Overall, eDNA metabarcoding analyses identified 52 fish genus/species (47 Actinopterygii and 5 Elasmobranchii) (Table S2), being the European anchovy (*Engraulis encrasicolus*) the most abundant one (Table S3), as in previous studies conducted in the same area (Fraija-Fernández et al. 2020). Almost half of species (25), accounting for 14% of the reads, were classified as deep-sea fish (Figure 2); among them, *Lophius piscatorius*, *Maurolicus muelleri*, *Xenodermichthys copei*, *Benthosema glaciale*, *Merluccius merluccius* and *Micromesistius poutassou* represented 95.9% of deep-sea fish reads, being widely spread along the water column, and detected at all depths (excepting *X. copei*, which was absent at 50m depth). Interestingly, also the most abundant epipelagic fish species were detected at all depths along the water column (Figure S1).

**Figure 2.**
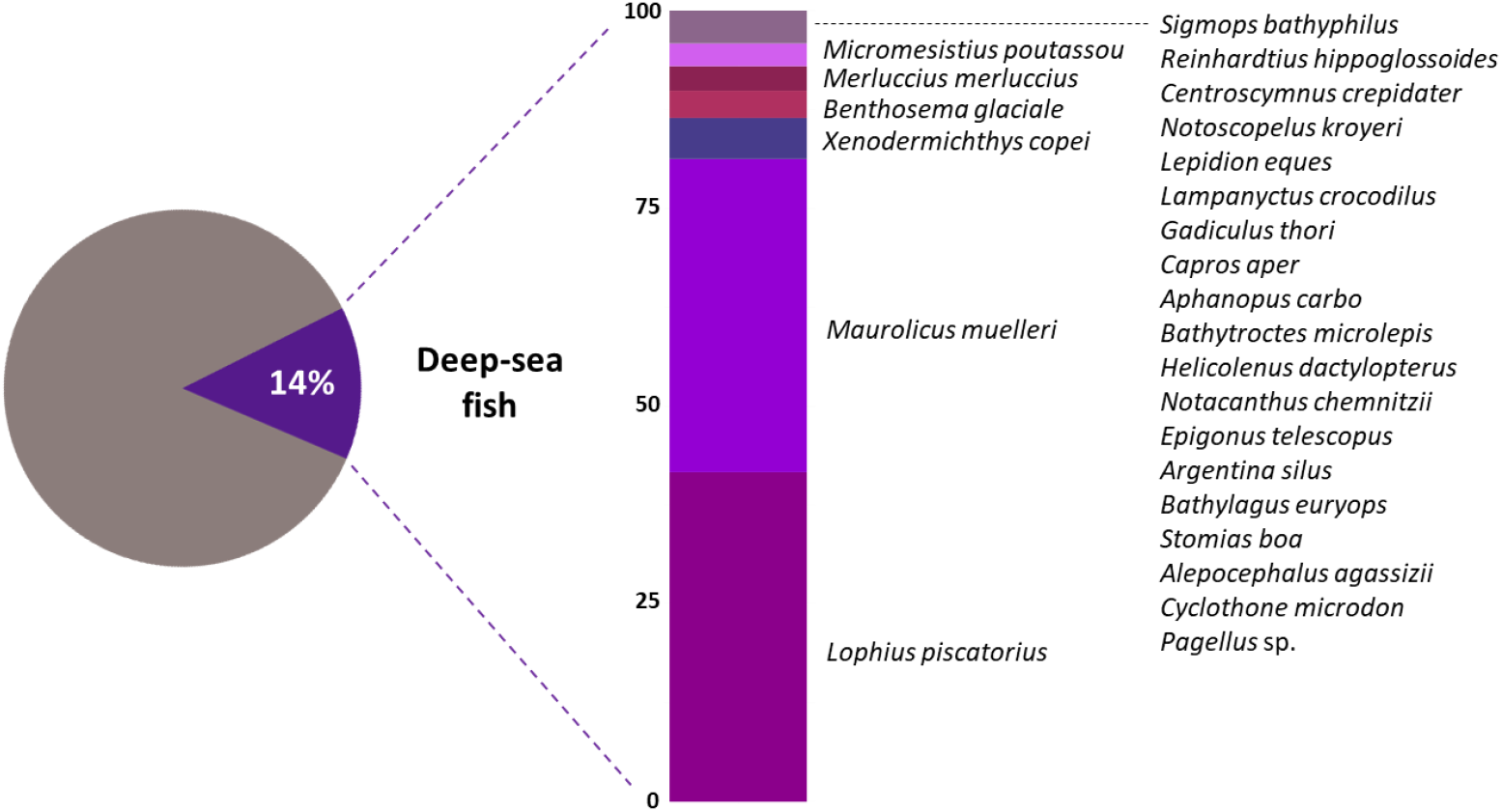
Proportion of reads belonging to deep-sea fish and within, relative abundance of each deep-sea fish species detected, ordered from the most (bottom) to the least (top) abundant.

### Deep-sea fish vertical structuring

An overall approach to fish vertical structuring assessed by eDNA metabarcoding showed an increase in deep-sea fish relative richness (proportion of deep-sea species related to total fish species) with depth (Figure S2). Deep-sea fish represented approximately 35% of richness in the epipelagic zone and over 50% in the mesopelagic and bathypelagic layers. In terms of abundance (percentage of deep-sea fish reads related to total fish reads per layer), deep-sea species accounted for less than 10% of relative abundance at the surface, and progressively increased to the mesopelagic zone, where it represented between 27 and 32% of fish abundance. The relative abundance of deep-sea species slightly dropped at > 1,000 m depth, accounting for about 17% of reads.

A more detailed analysis revealed that the vertical distribution patterns of deep-sea fish richness and abundance inferred from eDNA differed during day and night (Figure 3), which agrees with the diel vertical migratory behavior of several deep-sea species. In daylight-collected samples, deep-sea fish abundance peaked between 500 and 1,000 m depth (representing 33 and 39% of fish reads, respectively), but their DNA signal remarkably moved upward the water column at night, reaching over 75% of total fish reads in the epipelagic zone (50 and 200 m depth). Accordingly, daylight-collected samples showed an increase in deep-sea fish relative richness from 500 m depth, while the increase in deep-sea species richness moved up to 200 m depth in samples taken at night.

**Figure 3.**
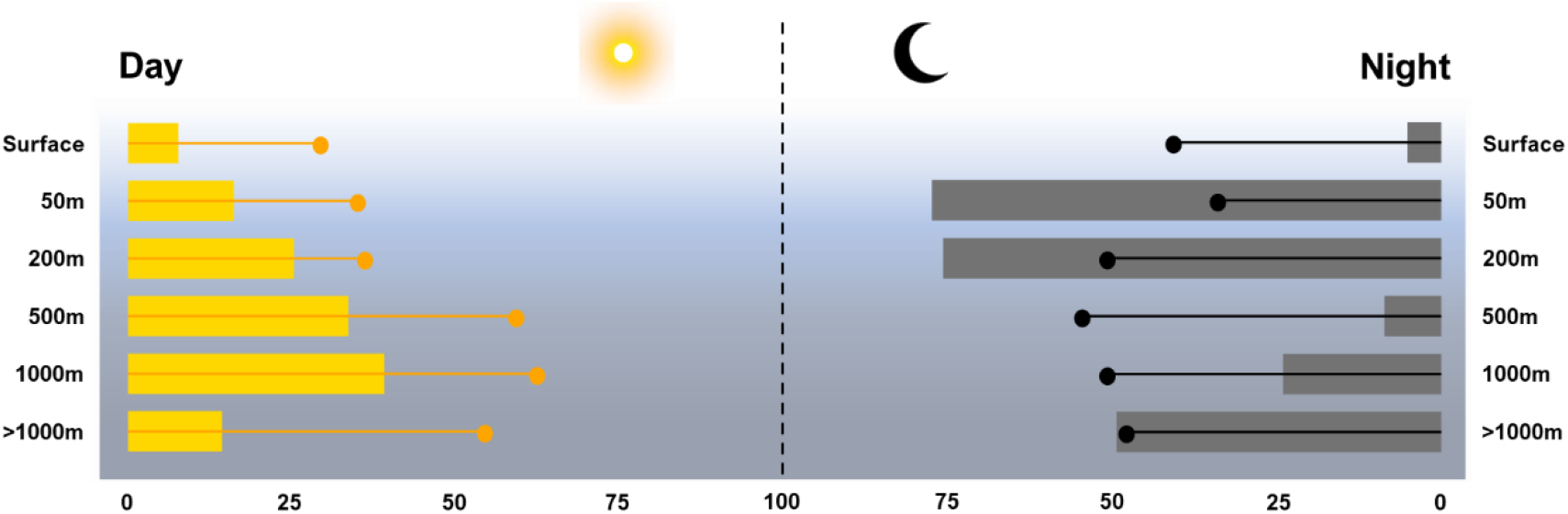
Relative abundance (bars, inferred from number of reads) and relative richness (dots) of deep-sea fish at each depth during day (left) and night (right).

### Fish species-specific DVM behavior

Noticeable, day-night variations along the water column emerged not only at the community level but also at the species level (Figure S3), indicating that eDNA metabarcoding data could be providing valuable insights on deep-sea fish species ecology and behavior. We observed four different patterns regarding diel vertical migratory behavior for the deep-sea species in our dataset (Figure 4): i) species without evidence of diel vertical migratory behavior appearing at all (or most) depths under study regardless of the time of sampling, i.e. day or night, such as *L. piscatorius*, ii) species appearing at all depths during both day and night but displaying evident DVM, e.g. *M. muelleri*, iii) species mainly observed from 200 m depth during daytime moving up to shallower waters at night, e.g. *B. glaciale*, and iv) species restricted to the mesopelagic and/or bathypelagic layers during both day and night, such as *Cyclotohone microdon*.

**Figure 4.**
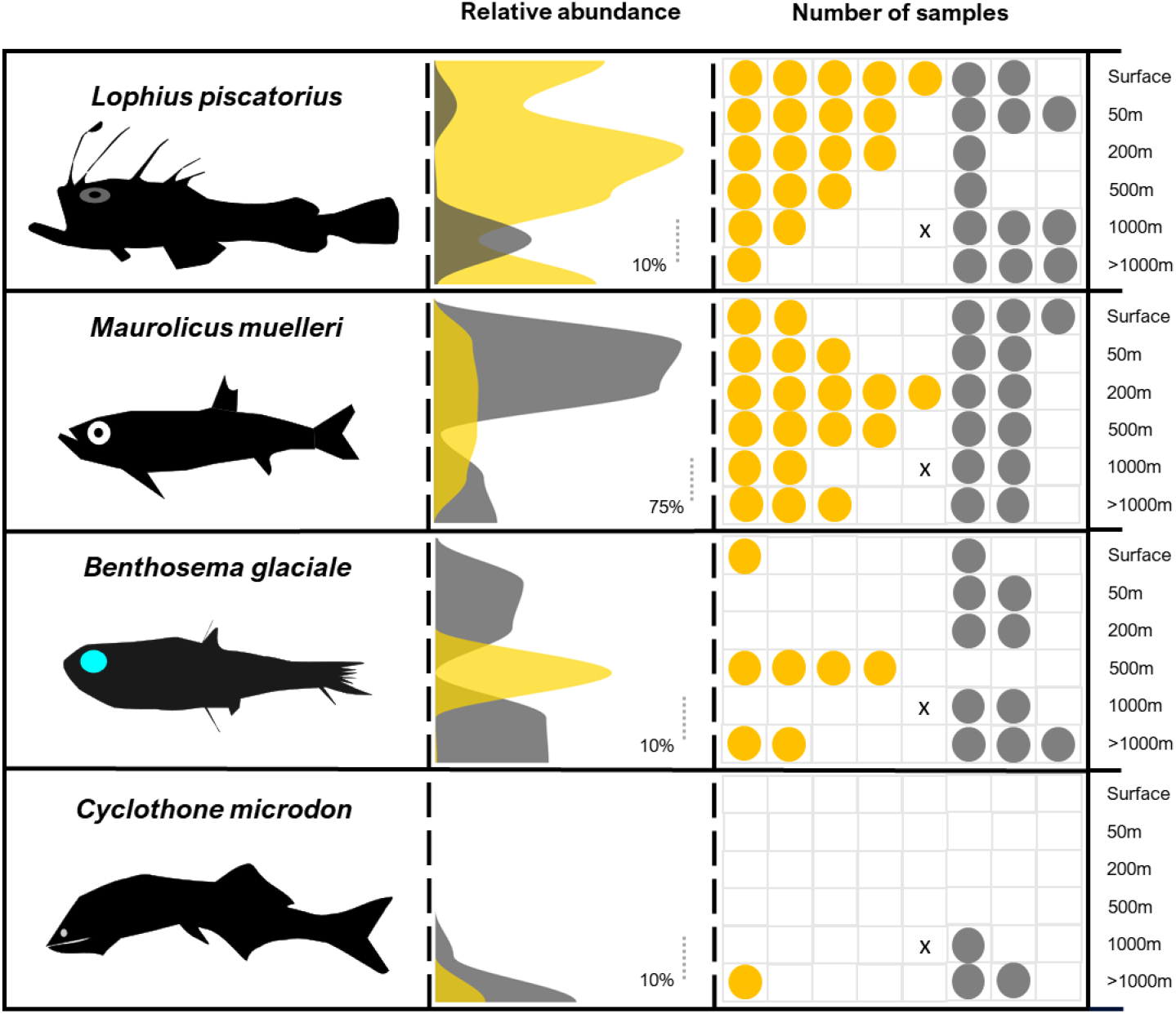
Vertical distribution of *Lophius piscatorius*, *Maurolicus muelleri*, *Benthosema glaciale* and *Cyclothone microdon* during day (gold) and night (gray) inferred from the percentage of reads (relative abundances) of each species at each depth studied, and the number of samples per depth at which these species were detected. A cross indicates that no data is available for that sample.

## Discussion

### Fish eDNA stratification in the ocean responds to fish vertical distribution

Despite the complexity of the vertical structuring of pelagic communities (Sutton 2013) and the call for studies focusing on the deep-sea ecosystem (Mengerink et al. 2014; St. John et al. 2016; Martin et al. 2020), to date just a few studies have used eDNA sampling to explore the deep-sea (Thomsen et al. 2016; Laroche et al. 2020; McClenaghan et al. 2020), and none has provided insights on the deep, vertical stratification of eDNA in the ocean. Here, we observed a consistent detection of DNA of the most abundant epipelagic fish species along the water column (e.g., European anchovy is detected down to >1,000 m depth), whereas detection of DNA from deep-sea fish was restricted to the upper depth at which they were assumed to occur and below. This implies that in the open ocean fish eDNA can be detected at the depth where it was released and deeper, but rarely at upper depths from which it was shed. These results highlight the potential of water column eDNA metabarcoding for the study of deep-sea organism ecology, such as diversity, spatial distribution, and their upper depth limit in the vertical gradient.

Our findings point to sinking-mediated vertical transport of fish eDNA along the oceanic water column. These findings are in line with the global sinking process of particulate organic matter in the ocean, according to which fish-released DNA is expected to sink from upper to deeper layers —either attached to or being part of bigger organic compounds such as feces, corpses, or scales, among others (Turner 2002). In this regard, previous studies have reported fish fecal pellets to present sinking rates ranging from hundreds to few thousands m·day^−1^ (Saba and Steinberg 2012), i.e., feces produced at the surface are expected to reach the mesopelagic zone in less than a day and, in some cases, even arrive to the bathypelagic realm in the same lapse of time. Hence, it may be deduced that the picture of fish community assessed by eDNA metabarcoding in the surface or at epipelagic depths is informative of the species present at the sampling point between the time of sampling and some hours before. The same principle could be also applied for deep-sea fish at least at the upper depth at which they are detected, although this assumption should be further confirmed by performing diurnal and nocturnal vertical profiles from the same sampling site. Noteworthy, following Preston et al. (2020), our findings open a lead to estimate fish species-specific contribution to the particulate carbon flux based on eDNA.

### eDNA can infer fish diel vertical migratory behavior

Probably, the most relevant, unexpected, and promising finding of the present study is the observation that eDNA metabarcoding is not only able to detect shifts at the community level derived by the DVM phenomena as suggested (Easson et al. 2020), but to reveal species specific diel vertical migratory patterns. Here, the most evident cases emerged for *Benthosema glaciale* and *Maurolicus muelleri*. During daylight, *B. glaciale* DNA was mainly found at 500 m depth and practically absent at upper layers, whereas at night, the molecular signal of *B. glaciale* moved upwards the water column, being widely detected at epipelagic depths. This clearly suggests DVM of *B. glaciale* from the deep-sea to the epipelagic zone at dark, in agreement with previous knowledge on the ecology of this species (Olivar et al. 2012). *M. muelleri* DNA appeared highly spread along the water column during both day and night, but exhibited remarkable abundance peaks during dark-time at 50 and 200 m depth. Although eDNA is to date not able to provide absolute biomass estimations, it is widely accepted that fish biomass and the amount of eDNA shed into the environment by fish are correlated (Klymus et al. 2015; Salter et al. 2019; Yates et al. 2020). Also, it has been demonstrated that eDNA shedding rates are influenced by diet, feeding activity —being up to 10-fold times higher in fed fishes than in starving ones— and life-cycle stage of the individuals (Klymus et al. 2015; Hansen et al. 2018). Thus, the noticeable high relative abundance of *M. muelleri* DNA at epipelagic depths during night may not only indicate a higher biomass but also elevated feeding activity, in line with the existing ecological knowledge on this species; *M. muelleri* is a very abundant sternoptychid in the Bay of Biscay, well-known to perform DVM in big, compact schools to feed close to the surface after the sunset (Sobradillo et al. 2019).

### Considerations for eDNA-based studies in oceanic vertical profiles

Notably, our eDNA-based data did not display vertical migratory patterns for some species known to migrate, e.g., *Lampanyctus crocodilus* or *Notoscopelus kroyeri*. Also, some species were detected only once or twice in the dataset (e.g., *Notacanthus chemnitzi*) and/or exclusively in diurnal vertical profiles (*Stomias boa*). Since most of these species presented very low abundances, the inconsistencies in their diel vertical migratory patterns and/or their scarce detection is probably explained by the relative low number of reads of these species in benefit of the highly abundant ones. In light of these findings, further studies should focus on enriching for deep-sea species or on increasing sequencing depths to obtain insightful information about low abundant species. Additionally, natural processes such as biological interactions or geological events can influence eDNA-derived inferences on fish vertical distribution. For instance, i) the predation of sharks, dolphins and other non-diel vertical migratory species on deep-sea fish, whose activity and high mobility could lead to detection of deep-sea fish DNA at shallow depths, ii) the effect of active convective cells, which can increase the vertical mixing of the water column (and therefore of fish eDNA) up to 700 m depth (Backhaus et al. 2003), and iii) the release of accumulated DNA in marine sediments —where DNA is more concentrated and much longer conserved (Turner et al. 2015)— during submarine landslides (Canals et al. 2006).

Notwithstanding, applying eDNA metabarcoding still offers remarkable advantages for the study of deep-sea ecosystems: it surpasses the depth-related orographic limitations imposed to trawling, and provides i) a higher diversity detection capability (Fraija-Fernández et al. 2020; Stoeckle et al. 2020), facilitating the monitoring of threatened, invasive, elusive, and rare species (Jerde 2019; Bani et al. 2020) —for instance, we detected here 6 fish species considered at least as vulnerable by the International Union for Conservation of Nature (IUCN; Table S3)—, ii) a lower dependency on taxonomic expertise (Hansen et al. 2018), iii) a reduction in costs (Jerde 2019; Bani et al. 2020), and iv) a minimum habitat destruction and organism disturbance (Valentini et al. 2016; Bani et al. 2020).

### Outlook

Protecting and preserving the remote deep-sea ecosystem is crucial to secure the global biogeochemical functioning of the planet, but techniques to monitor this environment, which is the basis of the development of protective measures, are lacking (St. John et al. 2016; Martin et al. 2020). Our findings support eDNA-based monitoring as a ready-to-use approach to acquire insights on this underexplored yet threatened environment as they provide evidence that the routine eDNA sample collection could lead to a significant advance in the understanding of deep-sea organismal ecology, including diversity, vertical structuring, and species-specific diel vertical migratory behavior. The incorporation of eDNA sampling to fisheries surveys has been shown to be economically affordable (Stoeckle et al. 2020), and the collection of vertical profile samples should not imply any important delay or reschedule of onboard activities (taking vertical profile samples with a rosette down to 2,000 m depths takes between one and a half and two hours long). This extra-effort certainly constitutes a small price to pay in exchange of acquiring the urgently needed knowledge on deep-sea environments to establish the basis of a sustainable extraction of its resources.

## Acknowledgements

Authors are grateful to the crew of R/V Ramon Margalef for their support during filtering and collection of samples, and specially to Beatriz Beldarrain, Maite Cuesta, Deniz Kukul and Carlota Pérez for their support on filtering onboard. Thanks to Elisabete Bilbao for technical assistance and Cristina Claver for useful input. This project has been supported by the Department of Economic Development and Infrastructure of Basque Government (projects GENPES, ECOPES and IM18MPDH), by the Spanish Ministry of Science and Innovation (project EDAMAME with reference CTM2017-89500-R) and by the European Union’s Horizon 2020 research and innovation program (project SUMMER with grant agreement No. 817806).

## Notes

### Competing Interest Statement

The authors have declared no competing interest.

### Summary of Updates

Correction in page 10; Maurolicus muelleri is not a myctophid but a sternoptychid.

