## Supplementary Material for "Vertical stratification of environmental DNA in the open ocean captures ecological patterns and behavior of deep-sea fishes"

### Supplementary Information

#### *DNA extraction and amplicon library preparation*

DNA extractions were performed using the DNeasy® blood and tissue kit (Qiagen) following the modified protocol for DNA extraction from Sterivex filters without preservation buffer by Spens et al. (2017*)*. DNA concentration was measured with the Quant-iT dsDNA HS assay kit using a Qubit® 2.0 Fluorometer (Life Technologies, California, USA). DNA was amplified with the teleo_F/teleo_R primer pair (hereafter 'teleo'), targeting a 60-70 bp region-long of the mitochondrial 12S rRNA gene, combined with the human blocking primer teleo_blk (Valentini et al. 2016). PCR mixtures were prepared under the hood in the pre-PCR laboratory using dedicated micropipettes and disposable plastic ware that were previously decontaminated under the UV light, and all postamplification steps were carried out in the post-PCR laboratory. PCR amplifications were done in triplicate with a final volume of 20 µl including 10 µl of 2X Phusion Master Mix (ThermoScientific, Massachusetss, USA), 0.4 µl of each amplification primer (final concentration of 0.2 µM), 4 µl of teleo_blk (final concentration of 2 µM), 3.2 µl of MilliQ water and 2 µl of 10 ng/µl template DNA. The thermocycling profile for PCR amplification included 3 min at 98 °C; 40 cycles of 10 s at 98 °C, 30 s at 55 °C and 45 s at 72 °C, and finally, 10 min at 72 °C. PCR products were pooled and purified using AMPure XP beads (Beckman Coulter, California, USA) following manufacturer's instructions, and used as templates for the generation of 12 x 8 dual-indexed amplicons in the second PCR reaction following the '16S Metagenomic Sequence Library Preparation' protocol (Illumina, California, USA) using the Nextera XT Index Kit (Illumina, California, USA). Multiplexed PCR products were purified using the AMPure XP beads, quantified using Quant-iT dsDNA HS assay kit using a Qubit® 2.0 Fluorometer (Life Technologies, California, USA) and adjusted to 4 nM. 5 µl of each sample were pooled, checked for size and concentration using the Agilent 2100 bioanalyzer (Agilent Technologies, California, USA), sequenced using the 2 x 300 paired end protocol on the Illumina MiSeq platform (Illumina, California, USA) and demultiplexed based on their barcode sequences.

*Read pre-processing, reference databases and taxonomic assignment*

Quality of raw demultiplexed reads was verified with FASTQC (Andrews, 2010). Forward and reverse primers were removed with cutadapt (Martin, 2011) allowing a maximum error rate of 20%, discarding read pairs that do not contain the two primer sequences and retaining only those reads longer than 30 nucleotides. Paired reads with a minimum overlap of 20 nucleotides were merged using Pear (Zhang et al. 2014), and those pairs with average quality lower than 25 Phred score were removed using Trimmomatic (Bolger et al. 2014). Reads that: i) did not cover the teleo region, ii) were shorter than 40 nucleotides, and iii) contained ambiguous positions, were removed using mothur (Schloss et al. 2009), as well as potential chimeras, which were detected based on the UCHIME algorithm (Edgar et al. 2011) . Taxonomic assignment of unique reads was performed according to the naïve Bayesian classifier method form Wang et al. (2007) implemented in mothur, and only reads classified to species level were considered for further steps. We used two reference databases for taxonomic assignment, named global and local databases, as described in Fraija-Fernández et al. (2020), which taxonomy was forced to match the seven taxonomic levels of the World Register of Marine Species (WoRMS; Horton et al., 2018): Phylum, Subphylum, Class, Order, Family, Genus, Species. The global database contained the teleo region from all Chordata sequences available from GenBank and was used to detect unexpected species and potential contaminations. Taxonomic assignment using the global database confirmed that most reads belonged to fish (519,358). Only 5 of them (0.001%) belonged to humans, while the remaining were assigned to Aves (0.1%) or were not successfully assigned to any Chordata class (“unclassified”, 1.2%).  The local database was restricted to the fish species (including Myxini, Petromyzonti, Holocephali, Elasmobranchii, Sarcopterygii and

Actinopterygii) expected in the Northeast Atlantic and Mediterranean areas and was used to assess fish diversity inferred from the study samples.

**Supplementary Material**

**Table S1.** For each sample collected, station, depth (m), geographic coordinates (latitude and longitude, degrees in sexagesimal notation), sampling time (local hour, GMT +1) and filtered volume (ml). * indicates the sample that was not successfully amplified.

| Sample_ID | Station | Depth (m) | Latitude | Longitude | Hour (Day - Night) | Filtered volume (ml) |
|---|---|---|---|---|---|---|
| 18BIO003 | PV_37 | 5 | 43.8770 | -6.3343 | 12.30h (Day) | 5000 |
| 18BIO004 | PV_37 | 50 | 43.8770 | -6.3343 | 12.30h (Day) | 5000 |
| 18BIO005 | PV_37 | 200 | 43.8770 | -6.3343 | 12.30h (Day) | 5000 |
| 18BIO006 | PV_37 | 500 | 43.8770 | -6.3343 | 12.30h (Day) | 5000 |
| 18BIO007 | PV_37 | 1000 | 43.8770 | -6.3343 | 12.30h (Day) | 5000 |
| 18BIO008 | PV_37 | 1580 | 43.8770 | -6.3343 | 12.30h (Day) | 5000 |
| 18BIO009 | PV_54 | 4.4 | 43.7743 | -4.5980 | 05.52h (Night) | 4200 |
| 18BIO010 | PV_54 | 5 | 43.7743 | -4.5980 | 05.52h (Night) | 4500 |
| 18BIO011 | PV_54 | 50 | 43.7743 | -4.5980 | 05.52h (Night) | 5000 |
| 18BIO012 | PV_54 | 200 | 43.7743 | -4.5980 | 05.52h (Night) | 5000 |
| 18BIO013 | PV_54 | 500 | 43.7743 | -4.5980 | 05.52h (Night) | 5000 |
| 18BIO014 | PV_54 | 1000 | 43.7743 | -4.5980 | 05.52h (Night) | 5000 |
| 18BIO015 | PV_54 | 1590 | 43.7743 | -4.5980 | 05.52h (Night) | 5000 |
| 18BIO023 | PV_111 | 4.4 | 43.6003 | -2.6883 | 21.07h (Night) | 5000 |
| 18BIO024 | PV_111 | 5 | 43.6003 | -2.7197 | 21.07h (Night) | 5000 |
| 18BIO025 | PV_111 | 50 | 43.6003 | -2.7197 | 21.07h (Night) | 5000 |
| 18BIO026 | PV_111 | 200 | 43.6003 | -2.7197 | 21.07h (Night) | 5000 |
| 18BIO027 | PV_111 | 500 | 43.6003 | -2.7197 | 21.07h (Night) | 5000 |
| 18BIO028 | PV_111 | 1000 | 43.6003 | -2.7197 | 21.07h (Night) | 5000 |
| 18BIO029 | PV_111 | 1150 | 43.6003 | -2.7197 | 21.07h (Night) | 5000 |
| 18BIO032 | PV_136 | 5 | 43.6743 | -2.1722 | 11.50h (Day) | 5000 |
| 18BIO034 | PV_136 | 50 | 43.6743 | -2.1722 | 11.50h (Day) | 5000 |
| 18BIO035 | PV_136 | 200 | 43.6743 | -2.1722 | 11.50h (Day) | 5000 |
| 18BIO036 | PV_136 | 500 | 43.6743 | -2.1722 | 11.50h (Day) | 5000 |
| 18BIO037 | PV_136 | 1000 | 43.6743 | -2.1722 | 11.50h (Day) | 5000 |
| 18BIO038 | PV_136 | 1170 | 43.6743 | -2.1722 | 11.50h (Day) | 5000 |
| 18BIO056 | PV_236 | 5 | 44.6250 | -2.3498 | 00.10h (Night) | 3860 |
| 18BIO057 | PV_236 | 50 | 44.6250 | -2.3498 | 00.10h (Night) | 5000 |
| 18BIO058 | PV_236 | 200 | 44.6250 | -2.3498 | 00.10h (Night) | 5000 |
| 18BIO059 | PV_236 | 500 | 44.6250 | -2.3498 | 00.10h (Night) | 5000 |
| 18BIO060 | PV_236 | 1000 | 44.6250 | -2.3498 | 00.10h (Night) | 5000 |
| 18BIO061 | PV_236 | 1290 | 44.6250 | -2.3498 | 00.10h (Night) | 5000 |
| 18BIO075 | PV_402 | 4.4 | 45.6250 | -3.7065 | 13.03h (Day) | 2050 |
| 18BIO076 | PV_402 | 5 | 45.6250 | -3.7065 | 13.03h (Day) | 5000 |
| 18BIO077 | PV_402 | 50 | 45.6250 | -3.7065 | 13.03h (Day) | 4270 |
| 18BIO078 | PV_402 | 200 | 45.6250 | -3.7065 | 13.03h (Day) | 5000 |
| 18BIO079 | PV_402 | 500 | 45.6250 | -3.7065 | 13.03h (Day) | 5000 |
| 18BIO080 * | PV_402 | 1000 | 45.6250 | -3.7065 | 13.03h (Day) | 5000 |
| 18BIO081 | PV_402 | 2080 | 45.6250 | -3.7065 | 13.03h (Day) | 5000 |
| 18BIO088 | PV_555 | 4.4 | 46.6233 | -5.1267 | 19.43h (Day) | 4600 |
| 18BIO089 | PV_555 | 5 | 46.6233 | -5.1267 | 19.43h (Day) | 5000 |
| 18BIO090 | PV_555 | 50 | 46.6233 | -5.1267 | 19.43h (Day) | 5000 |
| 18BIO091 | PV_555 | 200 | 46.6233 | -5.1267 | 19.43h (Day) | 5000 |
| 18BIO092 | PV_555 | 500 | 46.6233 | -5.1267 | 19.43h (Day) | 5000 |
| 18BIO093 | PV_555 | 1000 | 46.6233 | -5.1267 | 19.43h (Day) | 5000 |
| 18BIO094 | PV_555 | 1830 | 46.6233 | -5.1267 | 19.43h (Day) | 5000 |
| 18BIO098 | PV_629 | 4.4 | 47.3505 | -6.4293 | 16.35h (Day) | 2110 |
| 18BIO099 | PV_629 | 5 | 47.3505 | -6.4293 | 16.35h (Day) | 3000 |
| 18BIO100 | PV_629 | 50 | 47.3505 | -6.4293 | 16.35h (Day) | 5000 |
| 18BIO101 | PV_629 | 200 | 47.3505 | -6.4293 | 16.35h (Day) | 5000 |
| 18BIO102 | PV_629 | 500 | 47.3505 | -6.4293 | 16.35h (Day) | 5000 |
| 18BIO103 | PV_629 | 1000 | 47.3505 | -6.4293 | 16.35h (Day) | 5000 |
| 18BIO104 | PV_629 | 1340 | 47.3505 | -6.4293 | 16.35h (Day) | 5000 |

**Table S2.** Overview of the number of samples analyzed, total number of reads, number of reads assigned to each fish class, and number of taxa (genus/species) obtained for each depth.

| Depth | Samples | Total teleo reads | Reads assigned to Actinopterygii (%) | Reads assigned to Elasmobranchii (%) | Unassigned reads (%) | Actinopterygii taxa | Elasmobranchii taxa |
|---|---|---|---|---|---|---|---|
| **All depths** | **52** | **526,393** | **99.8 (96.8)** | **0.05 (0.01)** | **0.18** | **47** | **5** |
| Surface | 13 | 329,199 | 99.9 (99.1) | 0.07 (0.01) | 0.01 | 23 | 2 |
| 50m | 8 | 85,592 | 99.9 (95.0) | 0.05 (0.05) | 0.002 | 19 | 1 |
| 200m | 8 | 44,351 | 99.8 (95.7) | 0 | 0.17 | 24 | 0 |
| 500m | 8 | 35,544 | 98.7 (89.7) | 0.01 (0.01) | 1.3 | 20 | 1 |
| 1000m | 7 | 9021 | 97.5 (80.9) | 0 | 2.5 | 22 | 0 |
| >1000m | 8 | 22,686 | 99.3 (90.6) | 0.04 (0.01) | 0.71 | 30 | 2 |

**Table S3.** List of species with their frequency of detection (number of stations and samples), their overall and at each depth relative abundance (percentage of number of reads over the total), and their status in the IUCN red list (LC: least concern; NT: near threatened; VU: vulnerable; EN: endangered; CR: critically endangered; iucnredlist.org, last accessed July 2020).

| Family | Species (* indicates deep-sea) | Occurrence | | Relative abundance | | | | | | | Status IUCN |
|---|---|---|---|---|---|---|---|---|---|---|---|
| | | Stations | Samples | Overall | Surface | 50 m | 200 m | 500 m | 1000 m | > 1000 m | |
| Alepocephalidae | *Alepocephalus agassizii* * | 5 | 5 | 0.10 | 0 | 0 | 0.002 | 0 | 4.11 | 0.96 | LC |
| | *Bathytroctes microlepis* * | 1 | 1 | 0.01 | 0 | 0 | 0 | 0 | 0 | 0.33 | LC |
| | *Xenodermichthys copei* * | 7 | 13 | 0.72 | 0.50 | 0 | 0.11 | 5.05 | 1.42 | 1.27 | LC |
| Anguillidae | *Anguilla anguilla* | 2 | 2 | 0.0004 | 0.0003 | 0.001 | 0 | 0 | 0 | 0 | CR |
| Argentinidae | *Argentina silus* * | 2 | 2 | 0.02 | 0.002 | 0 | 0 | 0 | 1.26 | 0 | - |
| Bathylagidae | *Bathylagus euryops* * | 1 | 1 | 0.03 | 0 | 0 | 0 | 0 | 0 | 0.79 | - |
| Caproidae | *Capros aper* * | 1 | 1 | 0.01 | 0 | 0 | 0 | 0.08 | 0 | 0 | LC |
| Carangidae | *Trachurus trachurus* | 4 | 4 | 0.001 | 0.001 | 0 | 0.002 | 0 | 0 | 0 | VU |
| Carcharhinidae | *Prionace glauca* | 2 | 2 | 0.0004 | 0.0003 | 0 | 0 | 0 | 0 | 0.005 | NT |
| Cetorhinidae | *Cetorhinus maximus* | 1 | 1 | 0.004 | 0.01 | 0 | 0 | 0 | 0 | 0 | EN |
| Clupeidae | *Sardina pilchardus* | 8 | 43 | 4.95 | 3.50 | 0.30 | 20.76 | 11.27 | 11.73 | 1.58 | NT |
| | *Sprattus sprattus* | 8 | 28 | 10.07 | 7.04 | 23.34 | 5.86 | 11.78 | 0.10 | 15.15 | LC |
| Cyprinidae | *Phoxinus* spp. | 1 | 1 | 0.005 | 0 | 0 | 0 | 0 | 0 | 0.12 | - |
| Engraulidae | *Engraulis encrasicolus* | 8 | 47 | 66.99 | 77.67 | 51.88 | 37.72 | 44.24 | 42.86 | 61.43 | LC |
| Epigonidae | *Epigonus telescopus* * | 1 | 1 | 0.02 | 0 | 0 | 0 | 0 | 1.29 | 0 | LC |
| Gadidae | *Gadiculus thori* * | 1 | 1 | 0.002 | 0 | 0 | 0.02 | 0 | 0 | 0 | LC |
| | *Micromesistius poutassou* * | 8 | 22 | 0.40 | 0.12 | 1.25 | 0.85 | 0.13 | 2.12 | 0.35 | LC |
| | *Trisopterus* spp. | 1 | 1 | 0.02 | 0 | 0 | 0 | 0 | 1.25 | 0 | - |
| Gobiidae | *Aphia minuta* | 1 | 1 | 0.0002 | 0 | 0 | 0 | 0 | 0.01 | 0 | LC |
| | *Pomatoschistus* spp. | 1 | 1 | 0.002 | 0.002 | 0 | 0 | 0 | 0 | 0 | - |
| Gonostomatidae | *Cyclothone microdon* * | 4 | 4 | 0.10 | 0 | 0 | 0 | 0 | 0.32 | 2.39 | LC |
| | *Sigmops bathyphilus* * | 1 | 1 | 0.0002 | 0 | 0 | 0 | 0 | 0 | 0.005 | LC |
| Labridae | *Ctenolabrus rupestris* | 1 | 2 | 1.49 | 2.33 | 0.005 | 0 | 0 | 0 | 0 | LC |
| Lamnidae | *Lamna nasus* | 1 | 1 | 0.01 | 0 | 0.06 | 0 | 0 | 0 | 0 | CR |
| Lophiidae | *Lophius piscatorius* * | 8 | 32 | 5.77 | 6.07 | 3.34 | 8.77 | 5.36 | 2.73 | 5.98 | LC |
| Merlucciidae | *Merluccius merluccius* * | 8 | 25 | 0.43 | 0.13 | 0.002 | 1.69 | 0.75 | 5.33 | 2.06 | LC |
| Molidae | *Mola mola* | 6 | 9 | 0.18 | 0.28 | 0.001 | 0.002 | 0.003 | 0 | 0.10 | VU |
| Moridae | *Lepidion eques* * | 1 | 1 | 0.001 | 0 | 0 | 0 | 0 | 0.07 | 0 | LC |
| Moronidae | *Dicentrarchus labrax* | 5 | 9 | 0.004 | 0.003 | 0.01 | 0.002 | 0 | 0.01 | 0.01 | LC |
| Mugilidae | *Chelon* spp. | 6 | 12 | 0.01 | 0.003 | 0.02 | 0.01 | 0.01 | 0 | 0.08 | - |
| Myctophidae | *Benthosema glaciale* * | 8 | 17 | 0.48 | 0.01 | 0.36 | 0.44 | 5.25 | 1.90 | 0.52 | LC |
| | *Lampanyctus crocodilus* * | 3 | 4 | 0.001 | 0 | 0 | 0 | 0.02 | 0 | 0.005 | LC |
| | *Notoscopelus kroyeri* * | 4 | 4 | 0.001 | 0 | 0 | 0 | 0.01 | 0.04 | 0 | LC |
| Notacanthidae | *Notacanthus chemnitzii* * | 1 | 1 | 0.02 | 0 | 0 | 0 | 0 | 0 | 0.39 | LC |
| Pleuronectidae | *Reinhardtius hippoglossoides* * | 1 | 1 | 0.0002 | 0 | 0 | 0.002 | 0 | 0 | 0 | NT |
| Rajidae | *Leucoraja naevus* | 1 | 1 | 0.001 | 0 | 0 | 0 | 0.01 | 0 | 0 | LC |
| Salmonidae | *Salmo trutta* | 2 | 2 | 0.01 | 0 | 0 | 0.002 | 0 | 0 | 0.15 | LC |
| Scombridae | *Euthynnus alletteratus* | 1 | 2 | 0.001 | 0 | 0 | 0.005 | 0 | 0 | 0.01 | LC |
| | *Katsuwonus pelamis* | 1 | 1 | 0.001 | 0 | 0 | 0.01 | 0 | 0 | 0 | LC |
| | *Scomber* spp. | 8 | 37 | 2.28 | 1.83 | 1.95 | 3.09 | 4.83 | 9.81 | 2.37 | - |
| | *Thunnus* spp. | 4 | 4 | 0.04 | 0.0003 | 0.01 | 0 | 0 | 0 | 0.91 | - |
| Scophthalmidae | *Scophthalmus maximus* | 4 | 4 | 0.04 | 0 | 0 | 0.13 | 0.44 | 0 | 0 | VU |
| Sebastidae | *Helicolenus dactylopterus* * | 1 | 1 | 0.02 | 0 | 0 | 0 | 0.25 | 0 | 0 | LC |
| Soleidae | *Solea solea* | 3 | 5 | 0.02 | 0.0003 | 0 | 0.002 | 0 | 1.58 | 0.005 | LC |
| Somniosidae | *Centroscymnus crepidater* * | 1 | 1 | 0.0004 | 0 | 0 | 0 | 0 | 0 | 0.01 | LC |
| Sparidae | *Diplodus sargus* | 8 | 13 | 0.01 | 0.002 | 0.004 | 0.002 | 0.18 | 0 | 0.01 | LC |
| | *Pagellus* spp. * | 6 | 14 | 0.20 | 0.25 | 0.01 | 0.15 | 0.18 | 0.30 | 0.30 | - |
| | *Pagrus major* | 2 | 2 | 0.01 | 0.01 | 0.04 | 0 | 0 | 0 | 0 | NT |
| Sternoptychidae | *Maurolicus muelleri* * | 8 | 32 | 5.48 | 0.23 | 17.41 | 20.37 | 9.84 | 11.70 | 1.92 | LC |
| Stomiidae | *Stomias boa boa* * | 3 | 4 | 0.04 | 0 | 0.01 | 0 | 0.33 | 0.07 | 0.42 | LC |
| Trichiuridae | *Aphanopus carbo* * | 1 | 1 | 0.01 | 0 | 0 | 0 | 0 | 0 | 0.15 | LC |
| Xiphiidae | *Xiphias gladius* | 1 | 1 | 0.01 | 0 | 0 | 0 | 0 | 0 | 0.22 | LC |

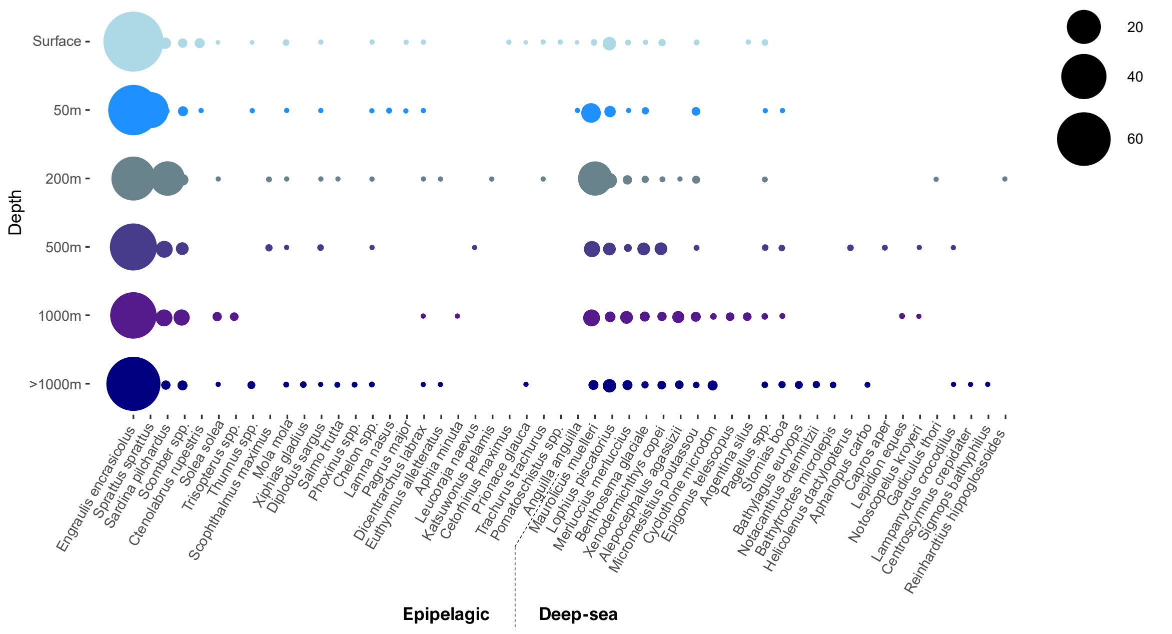


**Figure S1:** Vertical distribution of all fish species detected, where the size of the bubbles indicates the relative abundance in percentage of each species at a given depth.

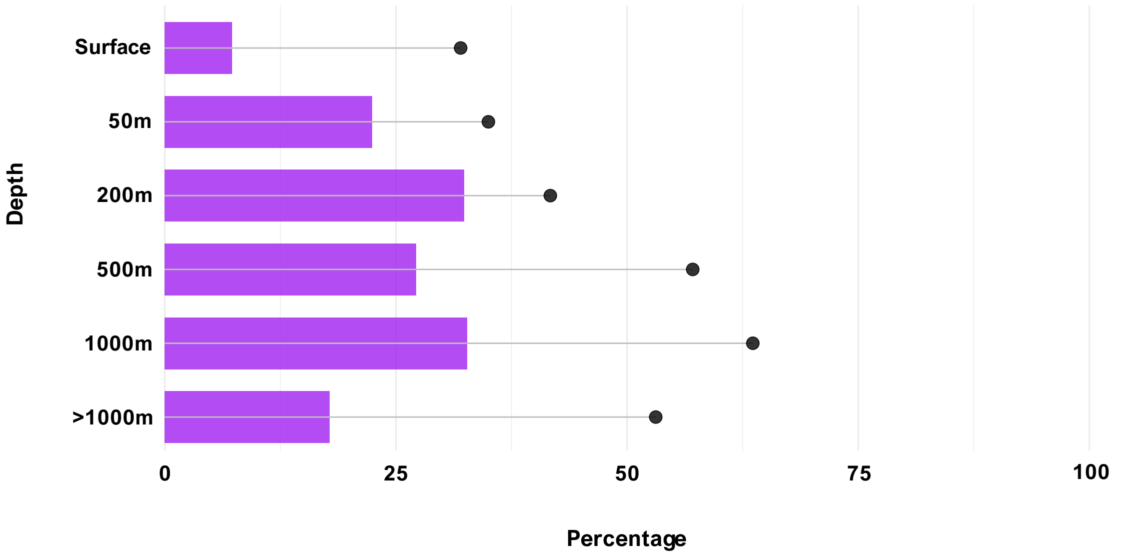


**Figure S2.** Relative richness (black dots) and abundance (horizontal bars, inferred from number of reads) of deep-sea fish with respect to whole fish community for each depth (including day and night vertical profiles).

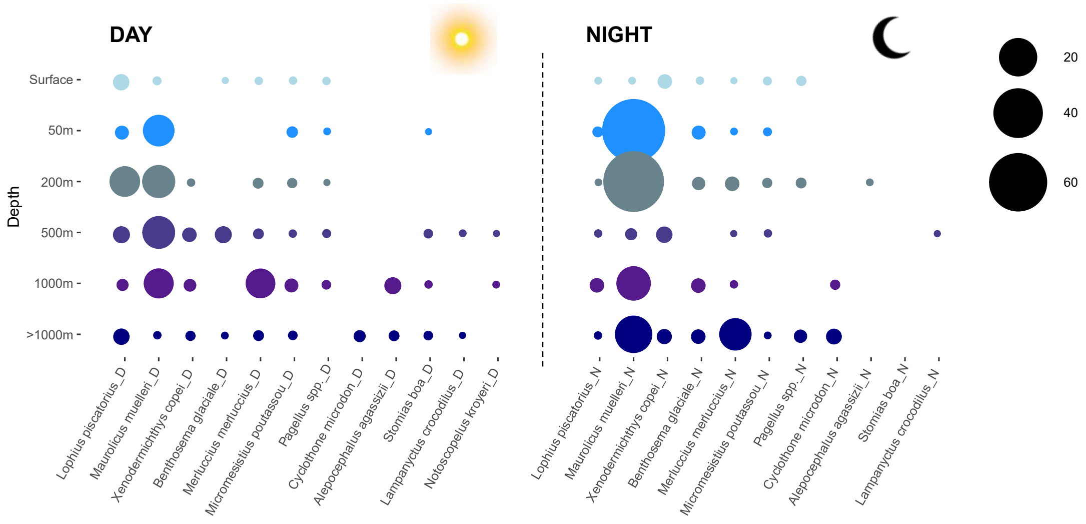


**Figure S3.** Vertical distribution of deep-sea fish species occurring in at least four of the samples, during day (left) and night (right). The size of the bubbles indicates the relative abundance of reads from each species at a given depth.
